# eMED-DNA: An *in silico* operating system for clinical medical data storage within the human genome

**DOI:** 10.1101/814830

**Authors:** Md. Jakaria, Kowshika Sarker, Mostofa Rafid Uddin, Md. Mohaiminul Islam, Trisha Das, Rameen Shakur, Md. Shamsuzzoha Bayzid

**Affiliations:** Bangladesh University of Engineering and Technology. Computer Science and Engineering, Dhaka, Bangladesh -1205.; The Koch institute for integrative cancer research at MIT. Massachusetts institute of technology 77 Massachusetts Ave, Cambridge, MA 02139, United States; Wellcome Sanger Institute. Wellcome Genome Campus. Hinxton. Cambridge CB10 1SA, UK

**Keywords:** Precision medicine, DNA storage, File management system, Genomic medicine, Translational technologies, Genome sequence, Non-coding regions, Next generation sequencing, Health Informatics

## Abstract

The propitious developments in molecular biology and next generation sequencing have enabled the possibility for DNA storage technologies. However, the full application and power of our genomic revolution have not been fully utilized in clinical medicine given a lack of transition from research to real world clinical practice. This has identified an increasing need for an operating system which allows for the transition from research to clinical use. We present eMED-DNA, an *in silico* operating system for archiving and managing all forms of electronic health records (EHRs) within one’s own copy of the sequenced genome to aid in the application and integration of genomic medicine within real world clinical practice. We incorporated an efficient and sophisticated *in*-DNA file management system for the lossless management of EHRs within a genome. This represents the first *in silico* integrative system which would bring closer the utopian ideal for integrating genotypic data with phenotypic clinical data for future medical practice.

## Introduction

The application of DNA as a digital information storage medium has long been considered a viable paradigm, but the real-world impact especially within the confines of actual clinical electronic health care record systems remains an enigma.^1–8^ The initial enthusiasm for such work was to enhance and better assimilate disparate health care information for each patient whilst integrating one’s underlying genomic data. Yet, the inherent problem with another form of data acquisition in health systems is the storage and assimilative properties of these often-large data files within the confines of clinical electronic medical systems and electronic health records (EHRs). Furthermore, the integration of genomic data coupled with multiple imaging modalities, consultation and prescription notes means this also poses a difficult task to formulate a robust, secure, accessible and transferable pathway to be applied in real world clinical settings.

In this regard, we demonstrate a simple, robust and informative computational clinical pipeline whereby genomic data across the genome can both be analyzed and simultaneously be integrated with one’s other clinical health records; hence providing a seamless archiving system for medical records and clinical genomic data. We have utilized a system in which a copy of the archived patient genome is used, and we particularly use the non-coding regions (introns and intergenic regions) of the human genome, to archive and store electronic health records as a proof of concept, whilst always maintaining the underlying archived genome sequence of the patient for future analysis. Moreover, our system can take any set of genomic locations, provided by a user, which is deemed not useful for a particular clinical context.

Our system is effective with any form of EHR, e.g., physician’s notes, lab reports, various image files which are typically stored in DICOM (Digital Imaging and Communications in Medicine) format. DICOM is the most universal and fundamental standard in digital medical imaging. It defines all the necessary file formats and network protocols to exchange the medical data. We selected computer files to be encoded as a proof of concept for practical DNA-storage, choosing a range of common medical formats to emphasize the ability of our system to store arbitrary digital information.

Given the variable spatio-temporal areas of non-coding regions across the human genome, which posed difficulty during the storage of files which span greater than the largest continual non-coding regions, we have innovated a sophisticated file management operating system that first encodes an EHR into DNA bases and then split it into chunks so that these chunks can be stored across various defined regions in the human genome. However, these regions will not affect analysis or interpretation of one’s genome given this is all done in a copied version, whilst archiving the original genome sequence. In particular, our contribution entails:

- An efficient *in*-DNA file management system (*i*DFMS).
- A novel technique for converting binary files to DNA sequences which is specially tailored for the binary data stream resulting from medical imaging files, but is suitable for any binary file. Our proposed encoding is more compact and better than the traditional binary to DNA mapping used in DNA storage.
- Analyzing and identifying various compression techniques appropriate for archiving EHRs.
- Introducing several concepts from operating systems (e.g., virtual memory, abstract memory space, dictionary for storing meta-data, etc.) to the domain of genome sequence, and DNA storage.
- Introducing efficient encoding (binary to DNA bases) and placement strategies for the *in*-DNA dictionary entries.

All these techniques are seamlessly combined into an integrated pipeline eMED-DNA for random and error free management of EHRs within a genome sequence. We believe that our system will facilitate the medical practitioners in managing and transferring both genomic and clinical phenotypic data across different medical and research institutes, and pave the way for meaningful integration of genotypes and phenotypes for precision medicine. Moreover, the new binary-to-DNA mapping strategy and *in*-DNA file management system, that we have introduced, will contribute towards the improvement of DNA storage technologies. We implemented eMED-DNA as a portable and cross-platform proof of concept software, which is publicly available at https://github.com/jakariamd/eMED-DNA. A comprehensive step-by-step video tutorial is available at https://jakariamd.github.io/eMED-DNA/.

## Results

### Archiving EHRs into DNA sequence

In this section, we provide a brief overview of eMED-DNA and some of its key components. We refer to the Methods section and supplementary materials SM1 for additional details of eMED-DNA and various algorithms used in it. Our pipeline (shown in Fig. 1) starts with compressing EHR components (e.g., physicians’ notes, DICOM files, bills, etc.) using appropriate compression techniques. For greater compression ratio, we explored various compression techniques for various types of EHR files and proposed customized compression techniques for DICOM images as these are the most space consuming components of EHRs. Next, binary bits of these compressed files are mapped to DNA base sequence {A, T, C, G}*. The naive and the simplest way to convert a binary sequence B = {0,1}* to a DNA base stream D = {A,T,C,G}* is to encode two bits of B into one bit of D. Suppose, one can encode 00 to A, 01 to T, 10 to C, and 11 to G. Another approach is phase-change encoding which takes the run-length of consecutive 0’s and 1’s into account, and was used in the Microvenus project.^10^ In run-length encoding, runs of 0’s and 1’s are stored as a single data value and count, rather than as the original run. We developed a new technique, especially tailored for DICOM files but suitable for any binary data, with improved performance compared with the trivial one which encodes two binary bits into a particular DNA base (see Methods section for full details of this encoding technique). Experiments on a CT (computerized tomography) scan with 262 DICOM files show that our method substantially improves upon the trivial mapping and saves more than one hundred and seventy thousand DNA bases for this CT scan (see Sec. 1.2 and Fig. S4 in supplementary materials SM1). The resultant DNA stream is subsequently compressed for further space efficiency.

**Figure 1:**
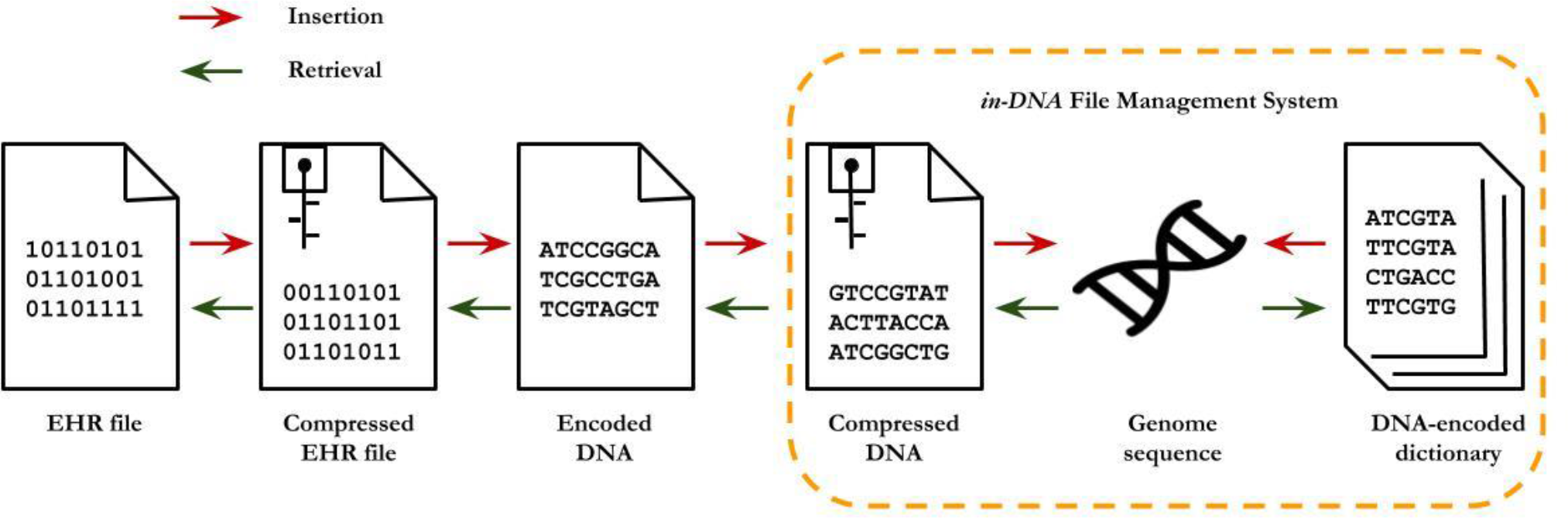
Overview of eMED-DNA. We first compress the EHRs with appropriate compression techniques. Next, the compressed binary files are mapped to DNA base sequence using our proposed binary to DNA base coversion technique. We further compress the DNA base stream, and finally we store the resultant DNA base stream within the non-coding regions (alternatively, the genomic regions provided by the user) of the patient’s genome. In addition to the EHRs, we encode the dictionary entries containing the meta-data into DNA bases and store them in the whole genome. The *in-*DNA file management system provides a random access and lossless architecture for archival and retrival of the data in a genome sequence.

Next, we store the resultant DNA bases in patient’s genome sequence. We divide the DNA streams into smaller chunks so we can spread them over the non-coding regions of the genome. We maintain an *in-*DNA dictionary for indexing the saved files and their positions in the genome so we can retrieve the stored files correctly. The dictionary itself is converted into DNA sequence and stored in the genome. It is consequently necessary to differentiate between DNA bases corresponding to EHRs from those of dictionary entries. Thus, we have to store and manage the dictionary efficiently so we do not lose any meta-data and at the same time we have to make sure it does not grow and overwrite the contents of an EHR which has already been written into the DNA sequence. For optimization and similar to the concept of the stack in random access memory (RAM), we write the dictionary in a reverse direction (relative to EHRs) starting from the end of the genome and progressing towards the front end (i.e., from right to left), whereas the EHR files are saved from the beginning of the genome (Fig. 3). See supplementary materials for full details.

eMED-DNA takes a human genome as an input. A reference human genome GRCH38 (Genome Reference Consortium Human Build 38), obtained from ENSEMBL, was used in our proof-of-concept. eMED-DNA also takes two other input files – one containing the locations (start and end markers) of the genes in chromosomes to identify the intergenic regions and the other containing the numbers of nucleotide bases in each chromosome. eMED-DNA can detect if any file has previously been saved in the given genome and shows the users a list of the saved files. A user can select, from the list, a filename to decode, so as to view the EHR or save it outside of the genome. eMED-DNA allows inserting new EHRs, as well as deleting the existing ones. The dictionary and the meta-data are updated accordingly after each operation.

### *in*-DNA File Management System (*i*DFMS)

In this section, we briefly discuss various key components of our proposed *i*DFMS. Non-coding regions are of various lengths and are distributed across the whole genome. As already mentioned, a stored EHR file may span several non-coding regions (fully or partially), and may span over more than one chromosome.

The deletion of a stored file from the DNA sequence will make the scenario even more complicated as it will free some regions in the DNA sequence which we have to mark for future use. To handle these challenges in a reasonably simpler way, we virtually assemble the non-coding locations from all chromosomes of a genome to form an abstract continuous space. We call this abstract space the *free genome space* (see Fig. 2).

**Figure 2:**
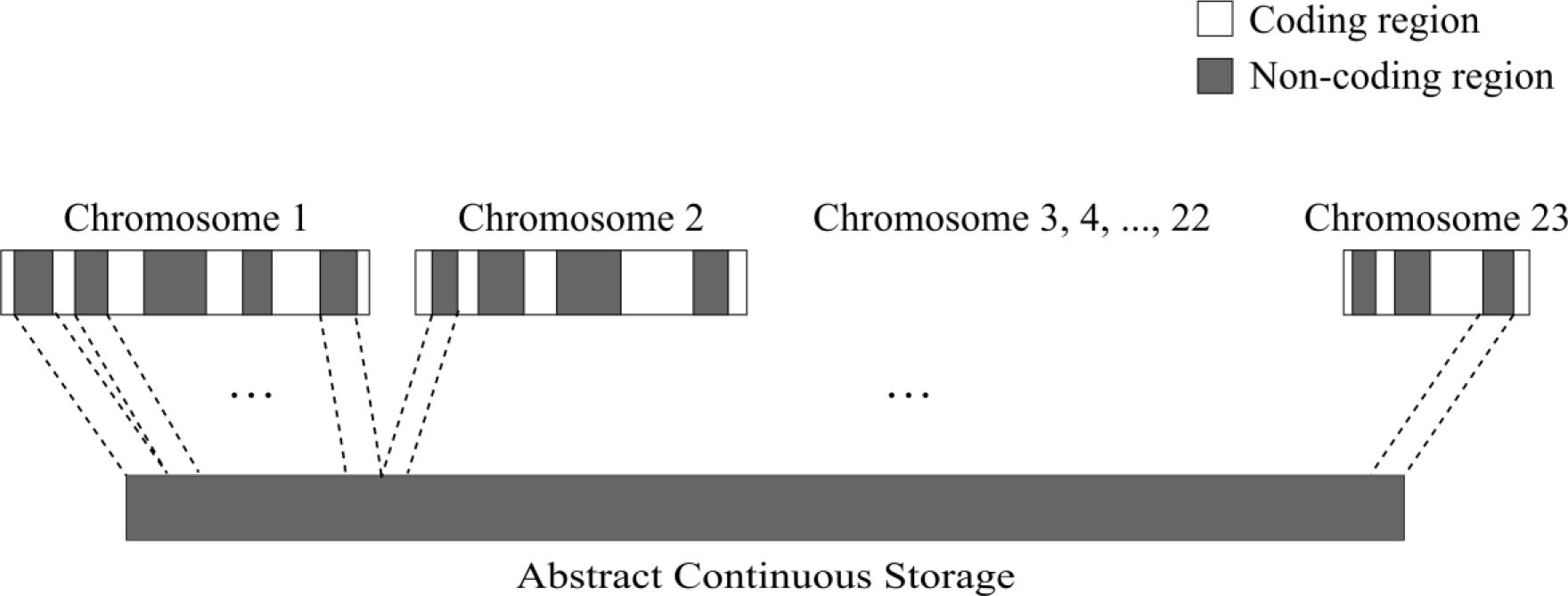
Abstract continuous space in a genome. eMED-DNA virtually assembles the non-coding regions (alternatively, it can assemble any set of genomic locations provided by the user to store EHRs) to an abstract continuous space.

We denote by a *user session* the time span, after entering a valid input set to the eMED-DNA system, for which the system remains actively open for that input set. At the beginning of every user session, our system constructs the corresponding abstract continuous storage space and creates a map between locations of abstract space and actual chromosome numbers and positions. In this way, a virtual abstract space (*free genome space*) is created in the beginning of a user session, resulting into simple and elegant file management without having to deal with complicated calculations for every file operations. While performing various file operations (insertion/deletion/retrieval), these abstract locations are converted to actual genomic positions appropriately.

### *in*-DNA Dictionary

We introduced an *in*-DNA dictionary in our *i*DFMS for storing various types of meta data. For each EHR file, we create an *entry* in the *in*-DNA dictionary which consists of the following four attributes: name of the file, type (DICOM or non-DICOM), transfer syntax and genomic locations containing the DNA bases of this particular file. When a genome is given as input, eMED-DNA can track the files (if any), which have already been saved in this genome. We have developed a new lossless mapping process to convert a dictionary to DNA base sequences (described in Methods) which is completely reversible so that it can decode the dictionary entries corresponding to the files that have already been stored in this genome in previous user sessions. We store the DNA bases resulting from the *in*-DNA dictionary in a direction which is opposite to the direction of storing DNA bases resulting from the EHRs (as shown in Figure 3). Please see Methods and Secs. 1.4 and 1.5 in supplementary materials SM1 for full details.

**Figure 3:**
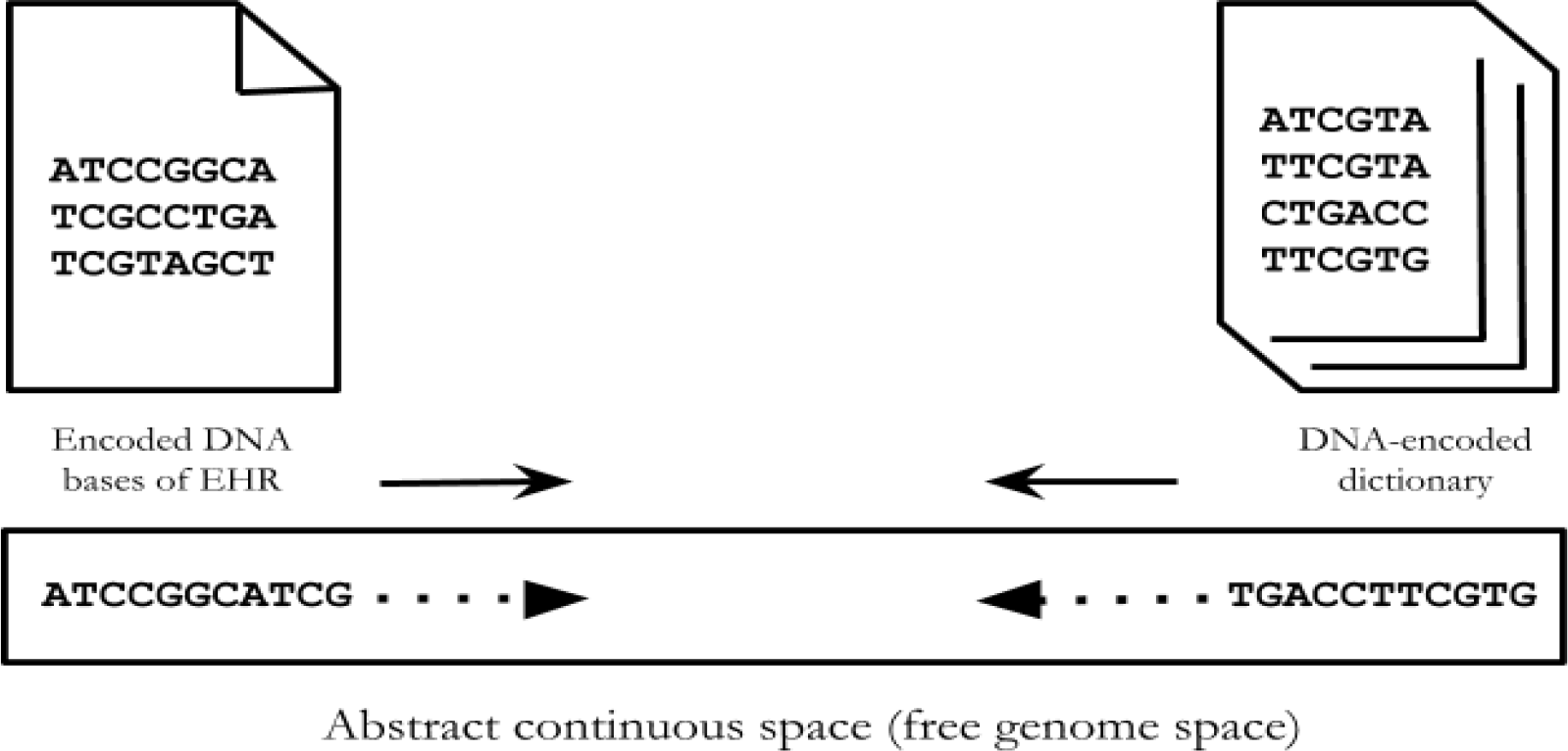
Placement of the EHR files and dictionary entries in a genome. We write the DNA bases resulting from EHRs from the beginning of the free genome space, as opposed to the *in-DNA* dictionary which starts from the end and grows towards the front end of a genome.

To emphasize the effectiveness of eMED-DNA in handling arbitrary digital information, we stored various types of EHRs using eMED-DNA and the summary statistics are provided in Table 1.

**Table 1:** Summary statistics of the various types of files stored using eMED-DNA. We choose various types of files to demonstrate the ability of eMED-DNA in handling any form of EHRs. These files are anonymized and available at https://git.io/fhIDE.

| File name | File type | File size<br>(bytes) | Number of bases<br>in the encoded<br>DNA stream | No of chunks<br>required |
| --- | --- | --- | --- | --- |
| MR-MONO2-8-16x-heart.dcm | Dicom (Magnetic Resonance Imaging) | 10,49,496 | 16,95,640 | 22 |
| US-PAL-8-10x-echo.dcm | Dicom (Echocardiogram) | 4,83,610 | 21,04,304 | 71 |
| Clinical note | Non-dicom<br>(physician's notes) | 2,264 | 5,060 | 2 |

Table 2 shows the distribution of the encoded DNA base streams over various chunks of non-coding regions for a single EHR file. See Sec. 1.7 in supplementary materials for additional results.

**Table 2.**
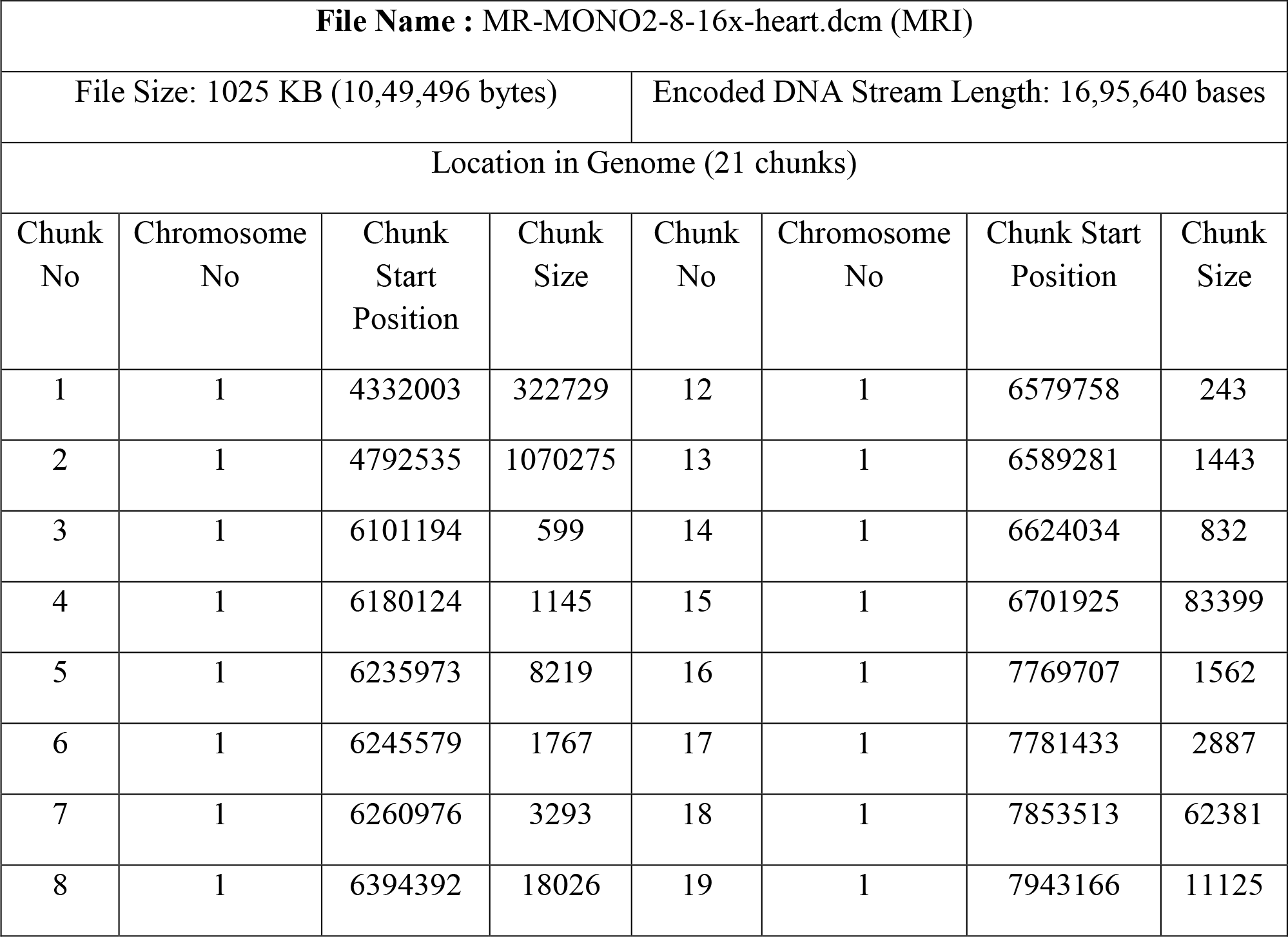
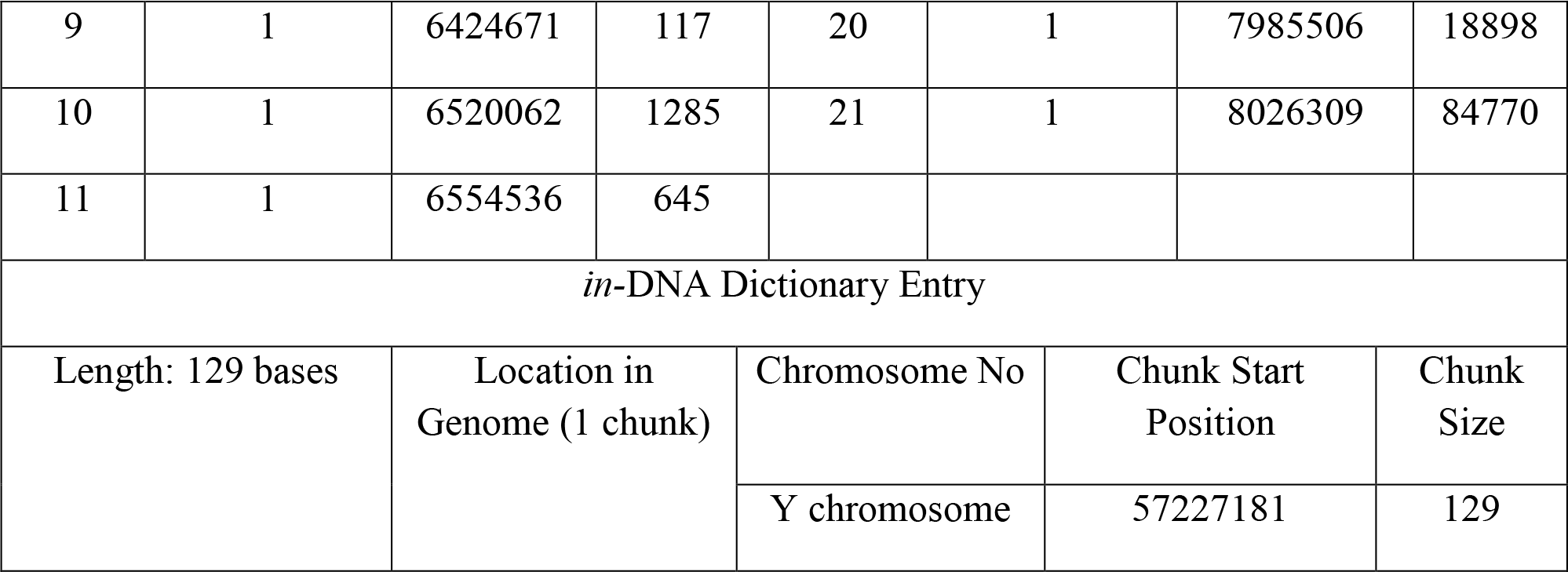
Distribution of the encoded DNA base streams of a sample DICOM file over various non-coding regions using eMED-DNA. This MRI file requires 21 chunks of intergenic regions of various sizes to fit the 16,95,640 encoded bases. The sample file is available at: https://git.io/fhIDq.

| File Name : MR-MONO2-8-16x-heart.dcm (MRI) |  |  |  |  |  |  |  |
| --- | --- | --- | --- | --- | --- | --- | --- |
| File Size: 1025 KB (10,49,496 bytes) |  |  |  | Encoded DNA Stream Length: 16,95,640 bases |  |  |  |
| Location in Genome (21 chunks) |  |  |  |  |  |  |  |
| Chunk No | Chromosome No | Chunk Start Position | Chunk Size | Chunk No | Chromosome No | Chunk Start Position | Chunk Size |
| 1 | 1 | 4332003 | 322729 | 12 | 1 | 6579758 | 243 |
| 2 | 1 | 4792535 | 1070275 | 13 | 1 | 6589281 | 1443 |
| 3 | 1 | 6101194 | 599 | 14 | 1 | 6624034 | 832 |
| 4 | 1 | 6180124 | 1145 | 15 | 1 | 6701925 | 83399 |
| 5 | 1 | 6235973 | 8219 | 16 | 1 | 7769707 | 1562 |
| 6 | 1 | 6245579 | 1767 | 17 | 1 | 7781433 | 2887 |
| 7 | 1 | 6260976 | 3293 | 18 | 1 | 7853513 | 62381 |
| 8 | 1 | 6394392 | 18026 | 19 | 1 | 7943166 | 11125 |
| 9 | 1 | 6424671 | 117 | 20 | 1 | 7985506 | 18898 |
| 10 | 1 | 6520062 | 1285 | 21 | 1 | 8026309 | 84770 |
| 11 | 1 | 6554536 | 645 |  |  |  |  |
| *in*-DNA Dictionary Entry |  |  |  |  |  |  |  |
| Length: 129 bases | Location in<br>Genome (1 chunk) | Chromosome No |  | Chunk Start<br>Position |  | Chunk<br>Size |  |
|  |  | Y chromosome |  | 57227181 |  | 129 |  |

## Discussion

Due to the advent of next generation sequencing within real world clinical medicine scenarios and the boom of the electronic health record systems to democratize personal health information, there has been an ongoing drive to integrate genomic data into clinical pathways. ^11–14^ The “All of Us” research program, formerly known as Precision Medicine Initiative, is expected to generate genomic data, in combination with electronic health records and participant-reported data, from approximately one million US residents with diverse backgrounds, so as to better risk stratify patients and enhance personalized diagnostics. ^15,16^

Precision medicine emphasizes the need for integrating genomic data and patient’s health records.^17,18^ However, including genomic sequences in EHRs has raised some complexities. Apart from security and ethical issues, there are practical challenges to integrate genomic data into electronic health records which include size and complexity of genetic test results, inadequate use of standards for clinical and genetic data, and limitations in EHR’s capacity to store and analyze genetic data.^19^ Furthermore, the plethora of large scale genomic studies has led to the discovery of many putative genes for better risk stratifying and enhancing diagnostic decision trees.^19–23^ Therefore, various large-scale initiatives have been taken for translating insights from genomics into clinical medicine.^11–14,23^ Hence, the efficient management, organization and protection of both genomic data and clinical records will be a greater necessity. eMED-DNA is a unique system for the delivery of genomic and clinical care in one operating system.

The eMED-DNA system not only converts generic file formats such as prescriptions, clinical lab results and physician reports, it also deals with clinical image files such as DICOM files and converts them into DNA sequences within the genome. This workflow can also be extended whereby these converted files can be synthesized into oligonucleotides and archived for a long period of time. Apart from the evident benefit in archiving, managing and transferring heterogeneous medical data across various medical and research institutes, this will have a direct impact in various retrospective medical studies. Although we did not perform any experimental studies to synthesize the genome sequence (obtained by eMED-DNA) into artificial DNA, existing DNA storage techniques can easily be coupled with eMED-DNA to facilitate artificial DNA synthesis. ^8–9^ Therefore, although we customized our encoding technique as an *in silico* system to avoid particular forms of repetitions, our system does not explicitly consider various issues associated with physical DNA storage, such as, repeats, homopolymers, small reads, etc. For example, Goldman et al. avoided repeats by translating the binary computer information into ternary number system and then encode the information into DNA bases.^5^ Yet, it is plausible that our *in silico* digital system can be part of a means to also identify and synthesise various EHR data for long term storage. Our proof-of-concept currently does not explicitly consider any privacy issue associated with genomic data sharing. However, the human genome can reveal sensitive information and is vulnerable to re-identifiability risk.^16, 24–30^ We believe a more sophisticated pipeline, which does not overwrite any region of the genome sequence such as ours, is more desirable as future research may reveal relevance of other regions which is currently considered as “junk” DNA. Indeed, recent research suggested that there may be relationships between introns and proteins, and long intergenic non-coding RNAs (lincRNAs) have been shown to have important functionalities.^31–32^ With a more sophisticated mapping system between EHRs and DNA bases, eMED-DNA would be able to retrieve the genomic regions, that it uses for storing EHRs, without having to keep an extra copy of the genome. (See Sec. 2 in supplementary materials for additional discussion).

The timing of this approach seems appropriate as obtaining genomic data are becoming easier and cheaper, DNA storage techniques are getting better which leads many to speculate on its near-term potential as a practical storage media^5^, population level genetic variation and understanding the genetic basis of diseases are getting significant attention from the research community^33–35^, and finally we are witnessing a rapid acceleration in the use of genomic information in patient care. We believe our proof-of-concept platform is the first of its kind and can greatly facilitate the future of precision medicine.

## Methods

### Encoding Binary files to DNA sequence

The naive and the simplest way to convert binary sequence B = {0,1}* to DNA base stream D = {A,T,C,G}* is to encode two bits of B into one bit of D. Suppose, one can encode 00 to A, 01 to T, 10 to C, and 11 to G. Another approach is phase-change encoding which takes the run-length of consecutive 0’s and 1’s into account, and was used in the Microvenus project.^36^ In run-length encoding, runs of 0’s and 1’s are stored as a single data value and count, rather than as the original run.

We developed a novel binary to DNA base conversion approach especially targeted for binary streams resulting from medical imaging files (DICOM). We observed the presence of long runs of 0’s in the binary streams of DICOM files which would be converted into long sequence of A’s in the usual encoding. Such repetitions are problematic for synthesizing, and are more likely to be misread by DNA-sequencing machines, leading to errors when reading the information back.^37^

Although homopolymers of short length does not make much problem in Illumina sequencing platforms, sequences of high or low GC content is difficult to sequence correctly.^1^ And higher frequency of A’s, resulting from long streams of 0’s, will decrease the GC content of the entire string.

Although we observed the presence of long runs of 0’s in the binary stream of DICOM files, the frequency of shorter runs of 0’s was much higher than the longer runs. For shorter runs of 0’s, the run-length based approaches give poorer compression than the naïve encoding. So we came up with an approach that does the naive encoding for shorter runs and run-length encoding for larger runs. We introduced a hyper parameter, which we call *‘shift’*, to serve this purpose. We do not take the long runs of 1’s into account as they are not usually prevalent in the binary streams of DICOMs due to their particular structures and grayscale values (see Figure S1 in supplementary materials SM1).

We read the binary bit stream by 2 bits at a time as the number of bits is always even (1 byte = 8 bits). We denote by *run-length* the length of a run of consecutive 0’s. If the *run-length* >= *shift* + *2*, we use run-length encoding; otherwise we use the naive encoding technique. Therefore, during the decoding step, we need to correctly differentiate the run-length encoded base streams from the base streams that have been encoded using the naive mapping. We use a prefix comprising a number of consecutive A’s equal to *shift*/2 +1 to mark the start of run-length encoding, and a single ‘A’ as a suffix to denote the end of the run. We store the run-length information between the prefix and the suffix by encoding the value of *difference.* We propose a new mapping technique to encode *difference* using DNA bases. Since ‘A’ is used as a suffix for the run-length information, the run-length information must be encoded with a ternary code that uses the remaining DNA symbols (C, G and T). One way to encode the difference would be converting *difference*/2 to ternary numbers (base 3), and finally mapping each of the base 3 digits (0, 1 and 2) to a different DNA base (T, C and G). However, by encoding *difference* using this approach, ternary numbers starting with 0 (e.g, 0 (T), 00 (TT), 000 (TTT), 010 (TCT), etc.) will never appear. In general, for *n* bits we should be able to use 3*n* permutations, but using simple base 3 conversion would only utilize 2*3^n−1^ permutations. Therefore, we have developed an efficient conversion technique (shown in Table S2 in supplementary materials SM1) which would utilize all possible permutations of the available three symbols, and thereby require fewer bits than the simple base 3 encoding described above.

Our technique is an incremental mapping between DNA bit sequence and the value of *difference*. It is designed in a way that enables us to represent run-lengths using minimum number of DNA bits possible. We developed a DNA base based number system accordingly. Suppose, ‘T’, ‘C’, and ‘G’ represent 1, 2 and 3, respectively. Then the numbers using only these characters are as follows: 1 (T), 2 (C), 3 (G), 11 (TT), 12 (TC), 13 (TG), 21 (CT), 22 (CC), and so on. The value of *difference* starts with 2 and is increased by 2 as we read two DNA bases at a time. Note that there is no DNA base mapping for the starting value (2) of *difference*. See supplementary materials SM1 for additional details.

### Converting the dictionary to DNA bases

For each stored file, we create an *entry* in the *in-*DNA dictionary which comprises the following attributes:

- Name: name of file (*f*).
- Type (*t*) (DICOM or non-DICOM): to select decompression procedure based on the type of the files.
- Transfer Syntax (*s*) (implicit or explicit): this is relevant only for DICOM files as lossless DICOM file decompression using j2k method requires transfer syntax.
- Genomic locations (*L*): a sorted list of all the chunks (with respect to the starting positions) which contain the sequence data of a particular file.

The dictionary of a genome itself is stored in the genome along with the actual files so that we do not have to manage separate files for meta data. After encoding a dictionary entry to a DNA base stream and storing it to the genome sequence, we need to be able to distinguish between various components (e.g, name, type, transfer syntax, genomic locations etc.) so we can read them back and decode the dictionary entry correctly. Therefore, we put some special markers in between these components. Note that these markers have to be encoded as DNA bases as well. DNA bases used to encode the markers should consequently be distinguishable from the encoded DNA base streams of the various dictionary components. Moreover, markers may be required between different dictionary entries to enable identification and reading of the individual entries correctly. Considering all these challenges, we have designed an efficient dictionary to DNA base stream conversion procedure which allows us to distinguish between individual dictionary entries and their components so we can decode them without any loss of information and ambiguity.

For a dictionary entry *e* with four components (*f*, *t*, *s*, *L*), we write them (starting from the tail end of the genome and progressing towards the front end) in the following order: *f*, *s*, *L* and *t*. We first describe the encoding technique of the file name (*f*) and the transfer syntax (*s*). The first 32 ASCII characters (out of 256) never appear in a file name as they are various control characters.

These 32 characters have 0’s in the three most significant positions in an 8-bit binary representation, which allows us to use “000” as a part of a marker. We add one more bit to incorporate the transfer syntax as well. That means, we use 0000 (AA) to denote the end of the file name as well as to denote the implicit transfer syntax, and use 0001 (AT) to denote the end of a file name and explicit transfer syntax. These two markers will not appear in the DNA base streams of the file names, and can be used to not only mark the end of the file name, but denote the transfer syntax as well.

We now describe the process for encoding *L* (genomic locations) and *t* (file type) of a dictionary entry. Let *L* be a set {*c*_1_, *c*_2_, *c*_3_, …, *c*_k_} of chunks in the genome sequence that make up a particular EHR. Here a chunk c_*i*_ is identified by its start position (*n*_1_) and size (*n*_2_). Note that *n*_1_ and *n*_2_ are integers and so they do not contain any character other than 0 ~ 9.

We represent each digit (0 ~ 9) by four bits and encode them into DNA base pairs as shown in Table S1 in SM1. We encode 00 by A, 01 by T, 10 by C and 11 by G. As already mentioned, the encoding to DNA bases is from right to left as as dictionary entries are written from the tail end of the genome, progressing towards the front. For a number *n* = *d*_k_*d*_k−1_…*d*_2_*d*_1_ with *k* digits, we start encoding and writing the digits into the genome sequence from the most significant bit (*d*_k_) and progress towards the least significant bit (*d*_1_). The corresponding DNA base pairs (as shown in Table 3) are appended from right to left. In this way, as the symbols are read back from right to left, they will be decoded in the appropriate order. For example, let *n* = 381. This will be encoded (starting from the most significant bit) as “TAACGA”. Therefore, while reading them back from right to left, we will have “AGCAAT” which will be decoded to 381 (considering two characters at a time) according to the example mapping.

**Table 3:** Mapping of numerical digits to DNA bases in dictionary to DNA base encoding. The base corresponding to the left 2 bits goes to the right. For example, for the digit 2, binary bits are 0010. Left 2 bits (00) map to ‘A’ and the right 2 bits (10) map to ‘C’. Now ‘A ‘goes to the right of ‘C’, i.e. final 2-length base string for digit 2 is ‘CA’.

| Digit | Binary | Base |
| --- | --- | --- |
| 0 | 0000 | AA |
| 1 | 0001 | TA |
| 2 | 0010 | CA |
| 3 | 0011 | GA |
| 4 | 0100 | AT |
| 5 | 0101 | TT |
| 6 | 0110 | CT |
| 7 | 0111 | GT |
| 8 | 1000 | AC |
| 9 | 1001 | TC |

There are 2^4^= 16 possible combinations of length four, of which only 10 combinations have been used in representing ten digits (see Table S3). The six remaining combinations: 1010 (CC), 1011 (GC), 1100 (AG), 1101 (TG), 1110 (CG), and 1111 (GG) may be used for marking the boundaries between various components and individual dictionary entries. Note that no digit in Table 3 starts with a ‘G’ (from right to left), and therefore we use ‘G’ as a separator to mark the end of a number *n*.

There are two combinations – 1010 (CC) and 1011 (GC) – that are unused in the encoded DNA base stream of *L*. We utilize these two combinations to mark the end of an entry *e* as well as to denote the file type *t* (DICOM and non-DICOM). For a DICOM file, CC may be used to mark the end of L, and GC may be used for non-DICOM files. Thus, by carefully designing the encoding techniques for various components of a dictionary entry and defining the order of encoding them, it is possible to distinguish between different components of a dictionary entry as well as the boundaries between individual entries without any ambiguity and loss of information. Figure 4 shows an example of a dictionary entry and the corresponding DNA base stream resulted from our encoding technique.

**Figure 4:**
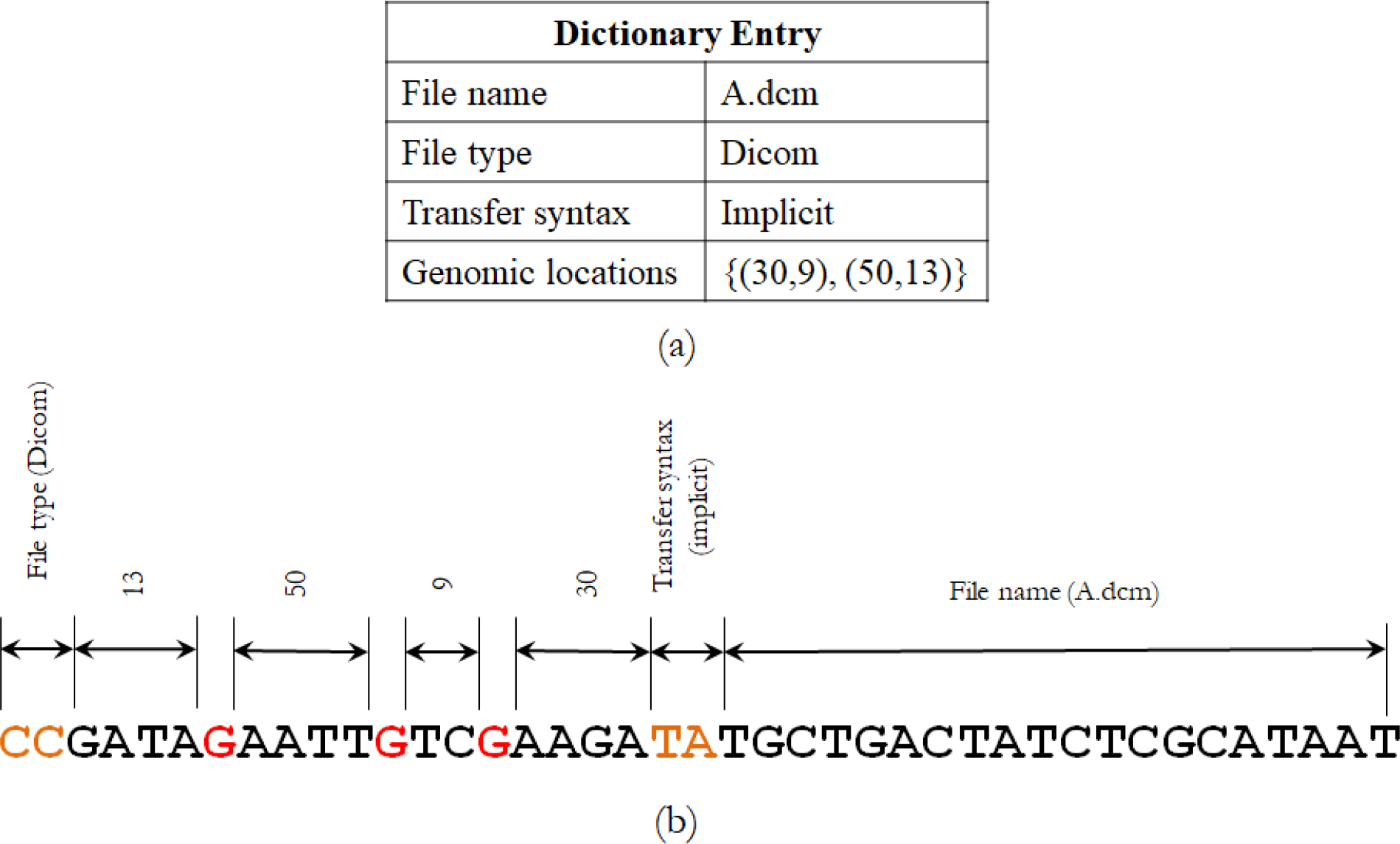
Encoding of a dictionary entry to DNA bases. (a) A dictionary entry with four components (name, type, transfer syntax and genomic locations). (b) Encoded DNA bases of this entry. We use 0001 (AT) to denote the end of the file name as well as the implicit transfer syntax, and CC is used to denote the end of the entry as well as the file type. In the DNA base stream of genomic locations (*L*), the G’s that are in the odd positions (marked in red) serve as markers to separate two numbers.

## Data Availability

Data supporting the findings of this study are available at https://git.io/fhIDE.

## Code Availability

The eMED-DNA software is freely available at https://github.com/jakariamd/eMED-DNA. The code is available on request from the authors.

## Author contributions

R. S and M.S.B conceived the study; M.S.B and R. S. supervised the study; M.S.B., M.J., K.S., M.R.U., M.M.I. and T.D. designed the eMED-DNA pipeline and developed the algorithms used in various components; M.J., K.S., M.R.U., M.M.I. and T.D. implemented eMED-DNA; M.J., K.S., M.R.U., T.D., M.M.I., M.S.B. and R.S. designed and conducted the experiments; M.S.B., R.S., M.J., K.S., M.R.U., T.D. and M.M.I. prepared the manuscript. The first five authors contributed equally and share the first-authorship of this paper.

## Supplementary Materials

These supplementary materials present additional details about various computational techniques and algorithms used in eMED-DNA (Section 1), and also present additional discussion (Section 2).

## 1. eMED-DNA

eMED-DNA is an *in silico* integrative pipeline which incorporates compression at different levels, a novel Binary-to-DNA base conversion technique, and an *in*-DNA file management system to manage the DNA base streams resulting from the electronic health records (EHRs) and the corresponding dictionary entries within ones genome sequence. We proposed customized algorithms and computational techniques for each of these key components of eMED-DNA.

### 1.1 Compressing DICOM files

In order to be able to accommodate large scale EHRs, we compress the medical records before encoding them as nucleotide bases. In particular, as DICOM files are the most space consuming and have special file formats, we proposed customized compression techniques suitable for DICOM files. We performed an extensive evaluation study to identify the techniques suitable for DICOM and finally, we customize the existing techniques so that they perform well on medical imaging files.

Compressing medical imaging files is challenging. The American College of Radiology (ACR) does not provide clear recommendation for compression.^1^ The US FDA does not permit compression storage or transmission of breast imaging. For other categories of image, we should carefully choose the type of image compression so that we do not loss any information and do not violate the DICOM standards. So, if there is a viable need for compression due to limited storage capacity, we should stick to lossless compression techniques.

There are many algorithms to compress binary data. But we only tried those which are recommended by DICOM standard.^2^ Current DICOM standard supports RLE^3^ (Run Length Encoding), Deflate^4^, JPEG^5^, JPEG LS^6^, and JPEG 2000^7^ compression techniques. Among these, JPEG is lossy while RLE, Deflate, JPEG LS and JPEG 2000 are lossless compression techniques. With an extensive evaluation study on various lossless compression techniques recommended by DICOM standards (results not shown), we decided to use JPEG 2000 for our first level of compression, i.e., compressing the binary data of the digital EHR file.

### 1.2 Compressed binary files to DNA sequence

We developed a new technique for encoding binary data to DNA base sequence (described in Methods section of this paper). In this section, we present additional details, examples and supporting experimental results.

**Figure S1:**
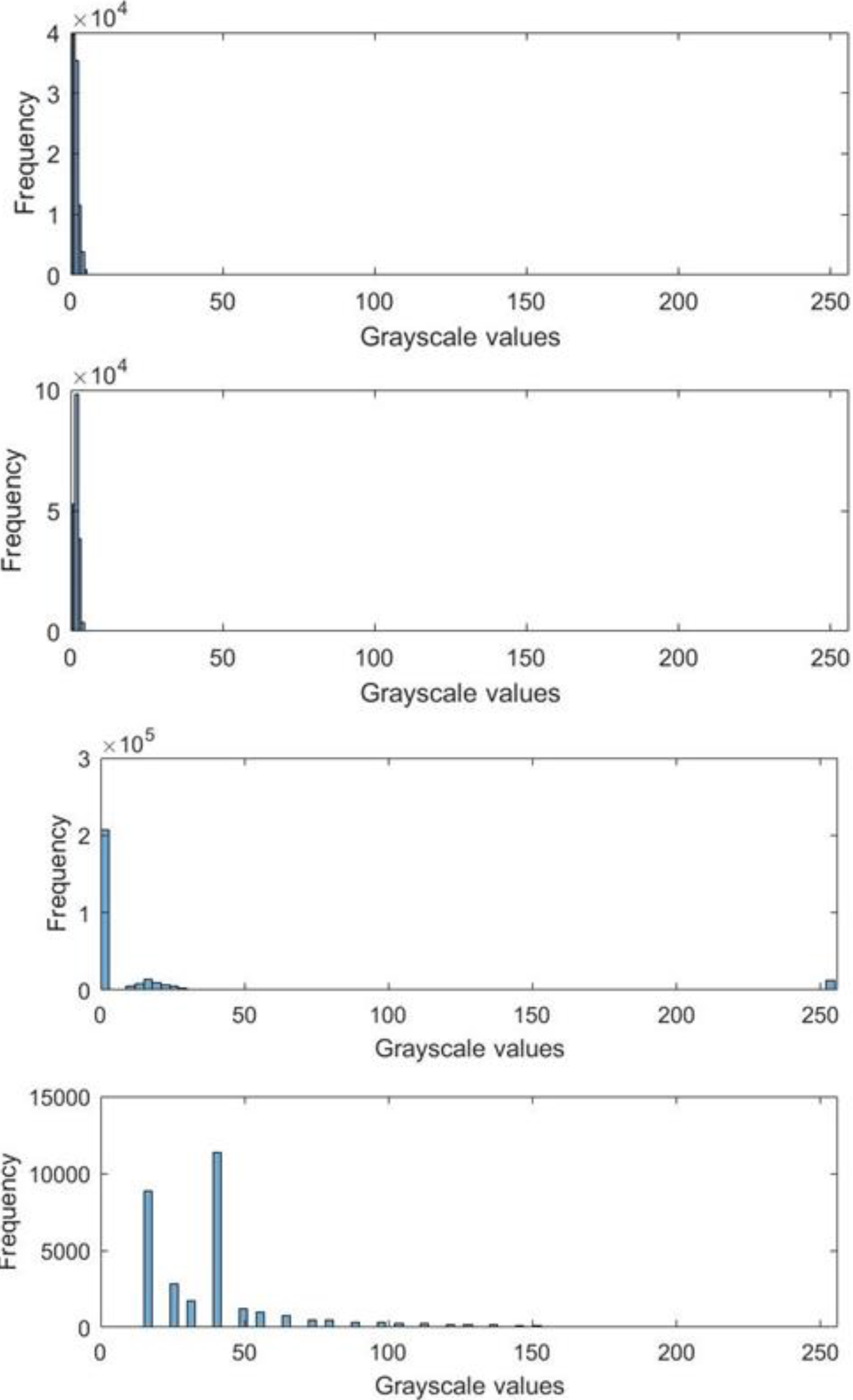
Histograms showing the frequencies of the pixel grayscale intensity values (0 ~ 255) for four sample compressed DICOM files. (top two plots are for RGB and the bottom two are for monochrome files). These files are available at: https://goo.gl/ruzozJ. All of these plots show right-skewed distribution. The frequencies of smaller values are much higher which reflects the presence of long runs of 0 in the binary stream of the Dicom files. This is due to the larger darker portions in DICOM files which have grayscale value of 0, i.e., RGB value of (0, 0, 0).

**Table S1:** A naive binary bits to DNA base ({0, 1}* → {A, T, C, G}*) encoding. Each of the four DNA bases can be represented by 2 binary bits. There are 4! = 24 such naive mappings. The table shows a mapping that we used for encoding compressed (non-DICOM) EHRs and the dictionary entries.

| Bits | Base |
| --- | --- |
| 00 | A |
| 01 | T |
| 10 | C |
| 11 | G |

**Table S2:** A novel run-length encoding of binary bits to DNA base sequence conversion. We show the mapping for values up to 76. This mapping allows us to utilize all possible 3^n^ permutations of three DNA bases.

| DNA bits | Run length – *shift* | DNA bits | Run length – *shift* | DNA bits | Run length – *shift* |
| --- | --- | --- | --- | --- | --- |
| - | 2 | TTT | 28 | CCC | 54 |
| T | 4 | TTC | 30 | CCG | 56 |
| C | 6 | TTG | 32 | CGT | 58 |
| G | 8 | TCT | 34 | CGC | 60 |
| TT | 10 | TCC | 36 | CGG | 62 |
| TC | 12 | TCG | 38 | GTT | 64 |
| TG | 14 | TGT | 40 | GTC | 66 |
| CT | 16 | TGC | 42 | GTG | 68 |
| CC | 18 | TGG | 44 | GCT | 70 |
| CG | 20 | CTT | 46 | GCC | 72 |
| GT | 22 | CTC | 48 | GCG | 74 |
| GC | 24 | CTG | 50 | GGT | 76 |
| GG | 26 | CCT | 52 | ... | ... |

By analyzing the mapping table (Table S2) we can see that if *difference* = *run-length* – *shift* is within the range [3^n^ +1, 3^n+1^ −1], we need *n* DNA bits to encode this *difference* value. Our encoding algorithm computes the value of *n* and converts (*difference* − 3^n^+1) /2 using a base-3 conversion and encodes the resulting ternary (base-3) number of *n* digits using a simple mapping as follows: {0 → T, 1 → C, 2 → G}. For example, let the value of *difference* be 22. This value (22) is within the range [3^2^ +1, 3^2+1^ −1], meaning two DNA symbols are required to encode *difference*. A base 3 conversion of (22 – (3^2^ +1))/2 = 6 is 20 [(6)_10_ = (20)_3_]. Finally, we map this to DNA bases (20 → GT) using the simple mapping mentioned above. Note that a direct base-3 conversion of this value (22) would require 3 DNA bits (CTG) as (22/2)_10_ = (11)_10_ = (102)_3_, whereas our method takes only 2 DNA bits.

This encoding technique not only offers higher storage capacity, it also allows us to design a lossless decoding algorithm. Our decoding technique reads the nucleotide base stream character by character. Unless the conditions for transitioning to run-length decoding are met, each character of the nucleotide sequence is simply mapped to two binary bits. However, in order to monitor for a transition marker, the decoding algorithm checks whether each ‘A’ is part of a run-length prefix by checking whether there are enough consecutive ‘A’s to be a prefix of run-length encoding. If a transition marker/prefix is identified, the DNA bases between the prefix and the suffix are read. Let the number of DNA bases in between the prefix and the suffix be *len.* The corresponding encoded *value* can be determined using the mapping (T → 0, C → 1, G → 2) followed by a base-3 to base-10 conversion. Next, the *difference* can be calculated as follows: *val*\*2 + (3^*n*^+1) = *difference.* Finally, a number of ‘0’s equal to *difference + shift* is added to the decoded binary bit stream. Suppose, the string between suffix and prefix is ‘GT’, then *len* equals 2 and *val* equals (20)_3_. After a base-10 conversion, *val* becomes 6 (since (20)_3_ = (6)_10_). So, a number of ‘0’s equal to 6*2 + *shift* + 3^2^ +1 or 22 + *shift* is added to the binary bit stream. The pseudo codes for encoding and decoding are shown in Fig. S2 and Fig. S3, respectively.

**Figure S2:**
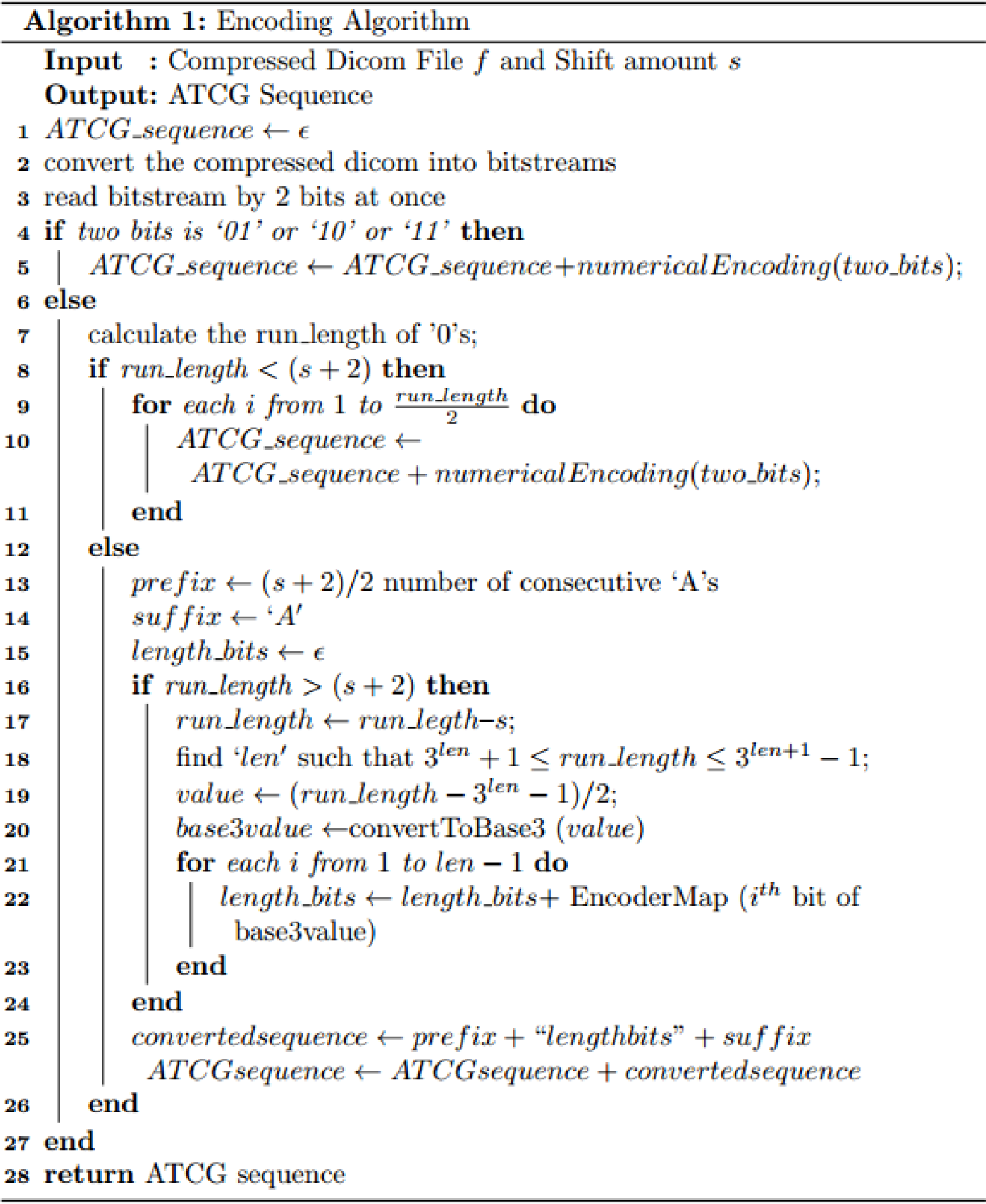
Binary sequence to DNA base stream encoding algorithm.

**Figure S3:**
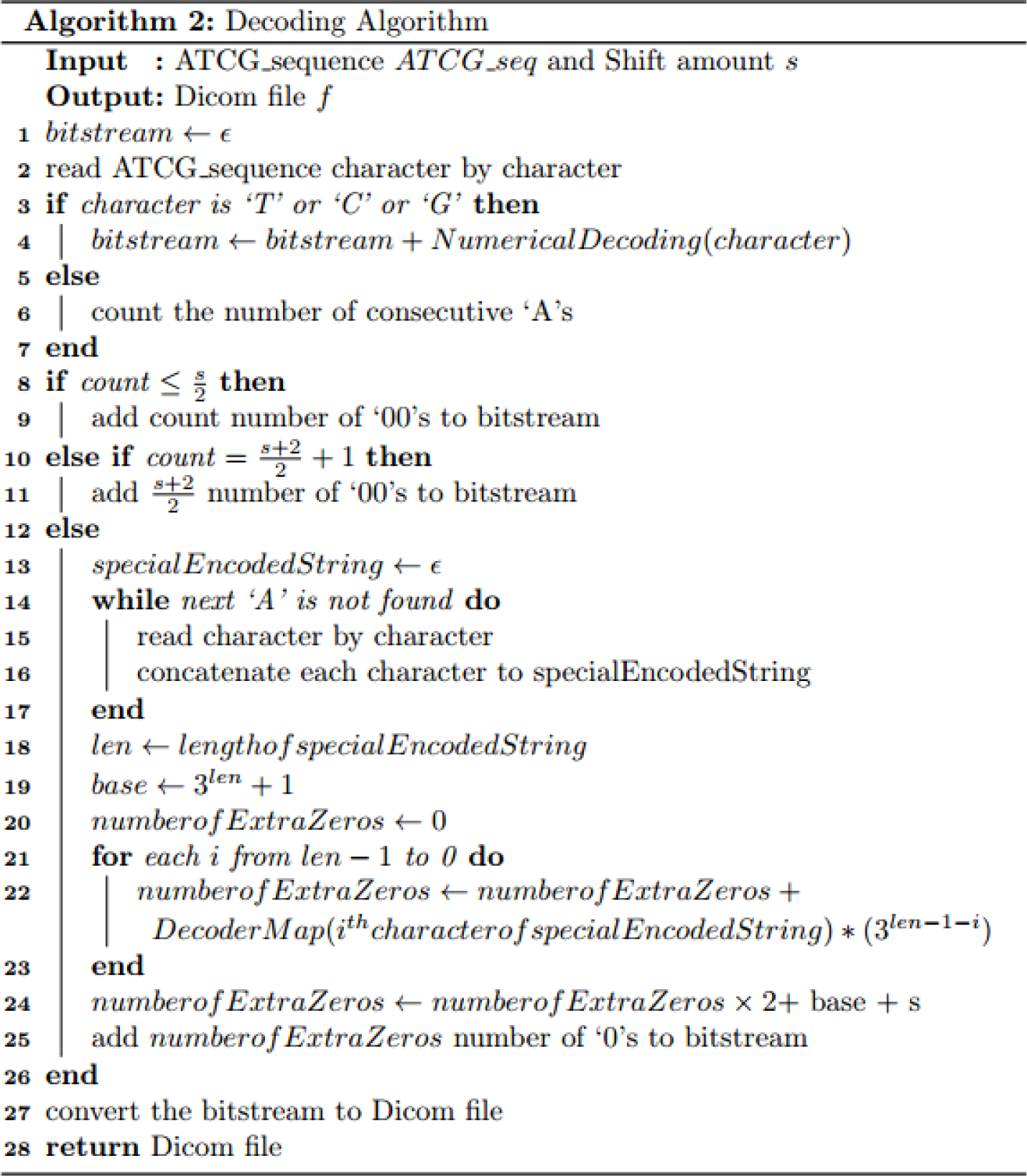
DNA base stream to binary sequence decoding algorithm.

Using this encoding and decoding techniques, we can achieve better compression ratio than the naive approach for most of the DICOMs that we tested. We evaluated the performance of our method on a CT (computerized tomography) scan of the skull consisting of 262 DICOM files and varied the values of the hyper parameter *‘shift’* (see Fig. S4). The average difference between our technique with *shift* = 16 and the naive method is around 650 DNA bases. Therefore, for the single CT scan we analyzed, the difference is around one hundred and seventy thousand DNA bases (262*650 = 1,70,300) which is clearly a substantial improvement. Thus our technique improves upon the naive mapping, especially when there is substantial amount of long runs of 0’s, which makes it suitable for DICOM files like CT scans. Further extensive evaluation on various types of DICOM files is required to better assess the performance of this encoding technique and identify a suitable value for the *shift* parameter.

**Figure S4:**
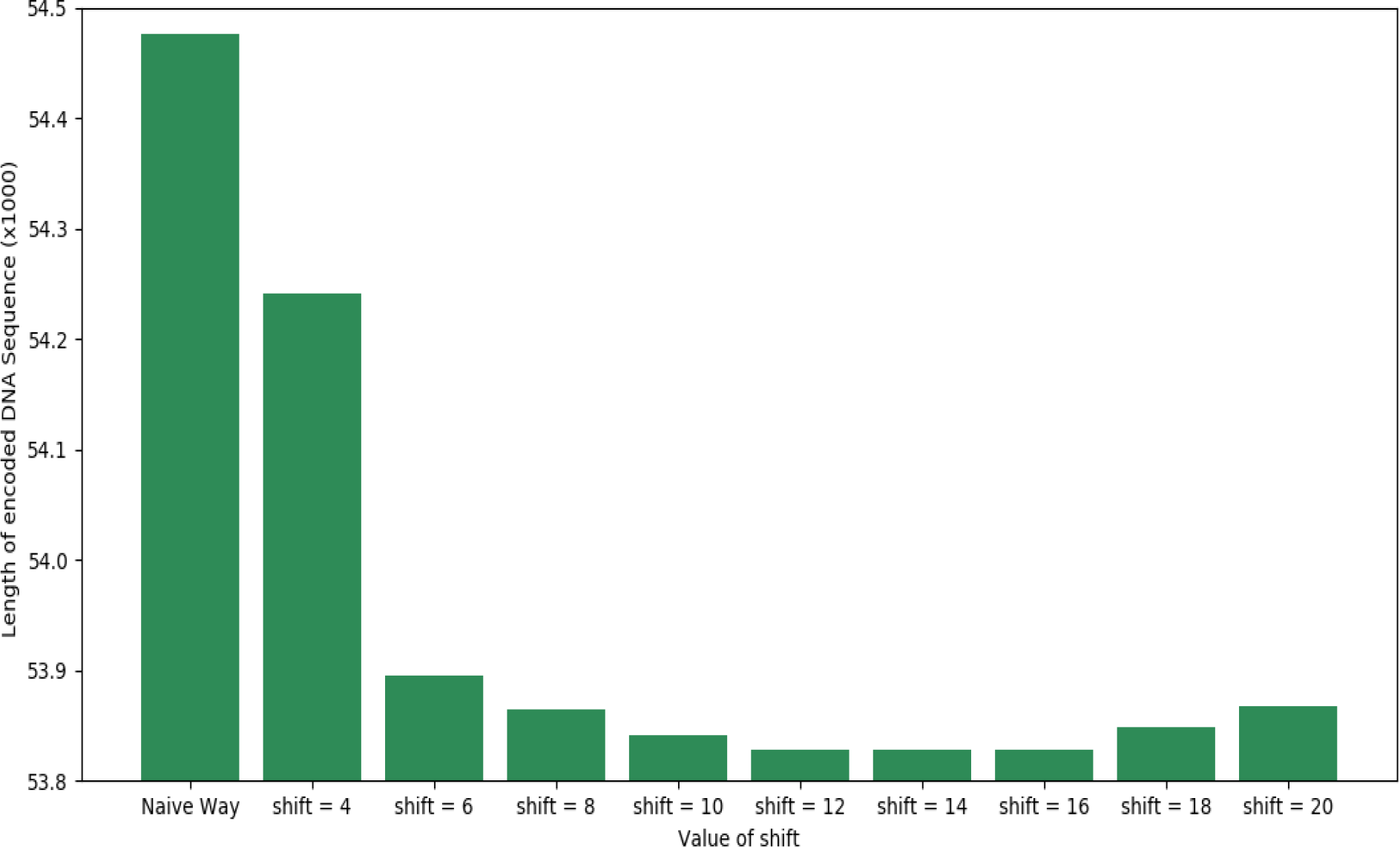
Comparison between our encoding technique (with various shift values) and the naive encoding (without shift parameter). We show the average lengths of the encoded DNA base streams on a single CT scan consisting of 262 DICOM files for the naive encoding and our technique with various values of shift (4, 6, 8, 10, 12, 14, 16, 18 and 20). The dataset contains 262 DICOM files in a CT scan of the skull collected from: https://wiki.cancerimagingarchive.net/display/Public/APOLLO-1-VA.

### 1.3 Compressing encoded DNA sequence

Once the mapping from binary streams to DNA base streams is done, we further compress the resulting DNA stream to achieve higher storage capacity within the DNA sequence. We performed an extensive evaluation study on various general purpose (Huffman^9^, Deflate^4^, RLE^3^, zip, 7zip^10^, bzip2^11^, gzip^12^, ANS^13^ etc.) as well as specialized compression techniques (BioCompress-I^14^, II^15^, DNACompress^16^, DNABit^17^, GenCompress^18^ etc.) for DNA sequence (results not shown). Considering the compression ratio, running time and ease of use of various techniques, we decided to use Deflate in eMED-DNA.

### 1.4 Placement of the dictionary in the genome sequence

As we have already mentioned, we start storing the dictionary (encoded as DNA stream) from the tail end of the genome. This is opposite to the direction of storing EHR files. If dictionary information were stored alongside the EHR files (from the same end of the genome), dictionary base streams would be interleaved with the EHR files. As dictionary entries produce extremely shorter base sequences than EHR files, the deletion of a file and its corresponding dictionary entry would potentially create small sized free chunks that would be available for storing future EHR files. The EHR files spread over these small free chunks would be fragmented, resulting in more meta information to store, which in turn results into larger dictionary entries. Storing the dictionary from the opposite end of a genome may address these problems.

#### Free List

Throughout an entire user session, we maintain a data structure called *free list* to keep track of the currently available free chunks (i.e., the genome space that are not currently being occupied with EHR files or the dictionary). Each user session starts with initializing the dictionary (by reading the *in*-DNA dictionary entries from previous user sessions) and constructing the free genome space accordingly.

### 1.5 File Operations

We now briefly describe various file operations within the genome sequence. Each user session of eMED-DNA starts with initializing the dictionary (by reading the *in*-DNA dictionary entries) and constructing the free list accordingly. eMED-DNA supports three basic file operations: insertion, retrieval and deletion.

#### Insertion

New EHR files may be added to a genome. eMED-DNA converts each input file to nucleotide base sequence according to the pipeline described above. For each EHR file, traversing the free list from front to end, the system makes a temporary list of chunks required to accommodate the encoded DNA base stream of that file. Next, a dictionary entry is formed with these locations and other meta data, and converted into DNA base stream. The DNA base stream of the dictionary entry is then inserted into the *in*-DNA dictionary and the DNA base streams of the EHR files are written into appropriate genomic locations, given that the genome has sufficient free space to accommodate these DNA base streams.

#### Retrieval

eMED-DNA can provide a list of the files that have already been stored in the genome sequence by acquiring necessary information from the *in*-DNA dictionary. eMED-DNA provides a graphical user interface so a user can select a file from this list to view or save a copy to the user’s computer. Once a file has been selected to view or save, eMED-DNA looks up the corresponding dictionary entry, obtain the genomic locations where this file has been stored and read the encoded DNA base sequences accordingly. Finally, eMED-DNA decodes the DNA base stream into a binary base stream by following a series of decompression and reverse mapping steps.

#### Deletion

With well-designed data structures, and EHR and dictionary placement techniques, removing a file from the genome becomes very straight forward. Removing an EHR file may comprise merely removing the corresponding entries in the *in*-DNA dictionary and updating the *free list* accordingly. All the entries after the deleted one are shifted by the size of the deleted entry, in order to mitigate fragmentation (similar to the “defragment” operation in an operation system).

### 1.7 Additional Results

Tables S3 and S4 show the distributions of the encoded DNA base streams over various chunks of non-coding regions for some representative EHR files.

**Table S3:** Distribution of the encoded DNA base stream of a sample DICOM file over various non-coding regions using eMED-DNA. This file requires 70 chunks of non-coding regions of various sizes to fit the 483,610 encoded bases. The sample file is available at https://git.io/fhIDq.

|  |  |  |  |  |  |  |  |
| --- | --- | --- | --- | --- | --- | --- | --- |
| File Name: US-PAL-8-10x-echo.dcm (Echocardiogram) |  |  |  |  |  |  |  |
| File Size: 473 KB (483610 bytes) |  |  |  | Encoded DNA Stream Length: 2104304 bases |  |  |  |
| Location in Genome (70 chunks) |  |  |  |  |  |  |  |
| Chunk No | Chromosome No | Chunk Start Position | Chunk Size | Chunk No | Chromosome No | Chunk Start Position | Chunk Size |
| 1 | 1 | 0 | 69091 | 36 | 1 | 1540454 | 1219 |
| 2 | 1 | 70009 | 112384 | 37 | 1 | 1574870 | 4886 |
| 3 | 1 | 184159 | 764 | 38 | 1 | 1580047 | 17964 |
| 4 | 1 | 200323 | 250417 | 39 | 1 | 1600097 | 15318 |
| 5 | 1 | 451679 | 234037 | 40 | 1 | 1630611 | 1484 |
| 6 | 1 | 686655 | 238225 | 41 | 1 | 1724325 | 513 |
| 7 | 1 | 959310 | 1277 | 42 | 1 | 1745993 | 5239 |
| 8 | 1 | 965716 | 781 | 43 | 1 | 1780458 | 4827 |
| 9 | 1 | 982094 | 16868 | 44 | 1 | 1891118 | 23709 |
| 10 | 1 | 1000173 | 965 | 45 | 1 | 1917297 | 293 |
| 11 | 1 | 1014542 | 5581 | 46 | 1 | 1919274 | 2677 |
| 12 | 1 | 1056119 | 14847 | 47 | 1 | 2003838 | 15486 |
| 13 | 1 | 1074308 | 7510 | 48 | 1 | 2030752 | 19718 |
| 14 | 1 | 1116362 | 57522 | 49 | 1 | 2212721 | 15974 |
| 15 | 1 | 1197936 | 5572 | 50 | 1 | 2310120 | 11133 |
| 16 | 1 | 1206692 | 4634 | 51 | 1 | 2391708 | 67 |
| 17 | 1 | 1214139 | 2769 | 52 | 1 | 2413798 | 12182 |
| 18 | 1 | 1232032 | 233 | 53 | 1 | 2505531 | 3002 |
| 19 | 1 | 1235042 | 7404 | 54 | 1 | 2526601 | 2144 |
| 20 | 1 | 1246723 | 7186 | 55 | 1 | 2530246 | 25393 |
| 21 | 1 | 1273886 | 6550 | 56 | 1 | 2565383 | 21108 |
| 22 | 1 | 1292030 | 346 | 57 | 1 | 2632991 | 2985 |
| 23 | 1 | 1324692 | 64 | 58 | 1 | 2801722 | 219761 |
| 24 | 1 | 1328898 | 2416 | 59 | 1 | 3022904 | 46264 |
| 25 | 1 | 1349351 | 3338 | 60 | 1 | 3438622 | 15804 |
| 26 | 1 | 1361778 | 11952 | 61 | 1 | 3481114 | 8806 |
| 27 | 1 | 1375496 | 10215 | 62 | 1 | 3611496 | 13506 |
| 28 | 1 | 1399339 | 2569 | 63 | 1 | 3630128 | 639 |
| 29 | 1 | 1407314 | 11106 | 64 | 1 | 3736202 | 16196 |
| 30 | 1 | 1421770 | 4358 | 65 | 1 | 3771646 | 1142 |
| 31 | 1 | 1427788 | 7073 | 66 | 1 | 3775983 | 2575 |
| 32 | 1 | 1442883 | 6806 | 67 | 1 | 3796505 | 15576 |
| 33 | 1 | 1470159 | 1610 | 68 | 1 | 3857215 | 52 |
| 34 | 1 | 1497849 | 14302 | 69 | 1 | 3885430 | 3695 |
| 35 | 1 | 1534688 | 486 | 70 | 1 | 3900294 | 431709 |
| *in*-DNA Dictionary Entry |  |  |  |  |  |  |  |
| Length: 105 bases |  | Location in Genome<br>(1 chunk) | Chromosome No |  | Chunk Start Position |  | Chunk Size |
|  |  |  | 24 |  | 57227310 |  | 105 |

**Table S4:** Distribution of the encoded DNA base stream of two clinical notes over various non-coding regions using eMED-DNA. Each of these two files requires two chunks (one for the note itself and the other one for the corresponding dictionary entry) of non-coding regions to fit the encoded bases. These notes were obtained from MIMIC-III database^19^.

|  |  |  |  |  |  |  |
| --- | --- | --- | --- | --- | --- | --- |
| File Name | NOTEEVENTS-00002.txt |  |  | NOTEEVENTS-00007.txt |  |  |
| File size | 2.21 KB (2264 bytes) |  |  | 1.77 KB (1813 bytes) |  |  |
| Encoded DNA Stream Length | 5060 bases |  |  | 4064 bases |  |  |
| Dictionary entry length | 107 bases |  |  | 107 bases |  |  |
| Location in genome | Chromosome No | Chunk Start Position | Chunk Size | Chromosome no. | Chunk Start Position | Chunk Size |
| EHR file (1 chunk) | 1 | 8111079 | 5060 | 1 | 8116139 | 4064 |
| Dictionary entry (1 chunk) | 24 | 57227074 | 107 | 24 | 57226967 | 107 |

## 2. Additional Discussion

This study presents a proof-of-concept for archiving medical records within the human genome sequences. This integration may last as long as thousands of years if preserved under appropriate environment using DNA storage, and thus will contribute towards the advanced research in personalized medicine and future healthcare system.^21,22^ In addition, our study makes significant contribution towards *in silico* DNA storage technology by introducing several novel techniques in coding theory and operating systems. Our proposed technique for binary-to-DNA conversion is especially tailored for DICOM files but can be used for any digital data.

eMED-DNA provides random access to the records stored in the genome sequence, and may offer the medical practitioners a one-stop solution for easy and efficient storage and management of both genotypes and phenotypes of the patients without having to deal with heterogeneous sources, types and standards of EHR data. eMED-DNA is expected to facilitate the transfer of medical records (along with the genome sequences) across various medical institutions as it can convert and store heterogeneous data into DNA bases and store them as simple computer files.

The ultimate goal of this framework is managing genotype-phenotype information of a patient’s lifetime in his DNA sequence. Therefore, we proposed compression techniques especially customized for medical records to accommodate the large amount of EHRs within the limited space available in a genome sequence. However, current compression ratio may not be sufficient to achieve the ultimate goal. Apart from developing advanced compression techniques, we can consider archiving large EHRs in a secured storage platform (for example, cloud storage) and store the link to the remote server along with necessary credentials in the genome sequence so that eMED-DNA can retrieve medical information from remote storage media.

This sort of initiative requires strict policies for privacy issues, and appropriate computational techniques for preventing any possible breach of privacy and information loss to unauthorized parties. Appropriate private key based cryptographic algorithms may be incorporated to eMED-DNA so that in case someone gets a genome sequence with encoded medical records, he will not be able to retrieve the information by using eMED-DNA unless he receives the private key from the original source. In addition to the encryption, strong identity authentication is required which guarantees that the sender and recipient of healthcare data, encoded in genomic sequence, are in fact who they claim to be.

In addition to storing and managing EHRs within ones genome sequence and thus facilitating genomic medicine, it would be interesting to add some other applications (relevant to genomic medicine) in eMED-DNA. For example, genome wide association study (GWAS) considers all the variants in the whole genome to see if any variant is associated to a particular trait or phenotype. The impact of GWAS in medical science could potentially be substantial as it contributes towards understanding the genetic factors contributing to variation in traits and diseases. Therefore, options for assisting GWAS by leveraging the genomic data and clinical phenotypes in eMED-DNA will be beneficial.

We believe eMED-DNA will have immediate positive impact in advanced medical research and DNA storage technologies. This will encourage the scientists to develop new architectures or enhance the ones presented in this study for better integration of phenotypic and genomic data. We believe this study will help the scientists to appreciate the need for integrating medical records with genomic data, and drive them towards utilizing this sort of frameworks for precision medicine.

